# A complex IRES at the 5’-UTR of a viral mRNA assembles a functional 48S complex via an uAUG intermediate

**DOI:** 10.1101/863761

**Authors:** Ritam Neupane, Vera P. Pisareva, Carlos F. Rodríguez, Andrey V. Pisarev, Israel S. Fernández

## Abstract

RNA viruses are pervasive entities in the biosphere with significant impact in human health and economically important livestock. As strict cellular parasites, RNA viruses abuse host resources, redirecting them towards viral replication needs. Taking control of the cellular apparatus for protein production is a requirement for virus progression and diverse strategies of cellular mimicry and/or ribosome hijacking evolved to ensure this control. Especially in complex eukaryotes, translation is a sophisticated process, with multiple mechanisms acting on ribosomes and mRNAs. The initiation stage of translation is specially regulated, involving multiple steps and the engagement of numerous initiation factors some of them of high complexity. The use of structured RNA sequences, called <u>I</u>nternal <u>R</u>ibosomal <u>E</u>ntry <u>S</u>ites (IRES), in viral RNAs is a widespread strategy for the exploitation of eukaryotic initiation. Using a combination of electron cryo-microscopy (cryo-EM) and reconstituted translation initiation assays with native components, we characterized how a novel IRES at the 5’-UTR of a viral RNA assembles a functional translation initiation complex via an uAUG intermediate, redirecting the cellular machinery for protein production towards viral messengers. The IRES features a novel extended, multi-domain architecture, circling the 40S head, leveraging ribosomal sites not previously described to be exploited by any IRES. The structures and accompanying functional data, illustrate the importance of 5’-UTR regions in translation regulation and underline the relevance of the untapped diversity of viral IRESs. Given the large number of new viruses metagenomic studies have uncovered, the quantity and diversity of mechanisms for translation hijacking encrypted in viral sequences may be seriously underestimated. Exploring this diversity could reveal novel avenues in the fight against these molecular pathogens.

## Introduction

Metagenomic studies of environmental samples have uncovered a fascinating diversity of viruses with a pervasive presence in the biosphere (*1, 2*). New viral clades have been discovered, showing an expanded presence compared with previously assumed distributions (*3*). Diversity is specially overwhelming in RNA viruses infecting animal hosts (*4*). Extensive gene shuffling, horizontal gene transfer events and host switching combined with co-divergency suggest a rich and complex evolutionary scenario within the animal virome, which remains largely unexplored (*1, 5*).

As strict cellular parasites, viruses rely on capturing cellular ribosomes to gain access to the host machinery for protein production (*6*). In eukaryotes, especially in animals, this machinery is complex and sophisticated, with large, multi-component protein factors assisting on the operation of eukaryotic ribosomes (*7*). Although complex, translation in eukaryotes conserves four main phases as its prokaryotic counterparts, namely: initiation, elongation, termination and recycling (*8*). Initiation is significantly expanded in eukaryotes, with two GTP-regulated steps required for the correct positioning of the first aminoacyl-tRNA responsible for setting up the correct reading frame on the messenger RNA (mRNA) (*9, 10*). In contrast to the streamlined mechanism of prokaryotic mRNA loading, docking of mRNAs to the eukaryotic small (40S) ribosomal subunit, requires several initiation factors (*10*). The mRNA molecules in eukaryotes are also covalently modified at their 5’ with a methylated guanine nucleotide called the “cap” and do not possess an equivalent of the Shine-Dalgarno prokaryotic sequence (*11, 12*).

Eukaryotic initiation starts when the 40S subunit together with initiation factors eIF1, eIF1A, eIF3, eIF5 and the <u>T</u>ernary <u>C</u>omplex (TC, eIF2/Met-tRNA_i_^Met^/GTP) form the 43S <u>P</u>re-Initiation <u>C</u>omplex (43S-PIC) which is competent for mRNA recruitment (*9*). Eukaryotic mRNAs are then docked to the 43S-PIC at their 5’ ends, forming the 48S complex (*13*). This complex is highly dynamic, able to move in the 5’ to 3’ direction along the mRNA in search of an AUG initiation codon in a favorable context, a process called “scanning” (*12*). Once the AUG codon is detected, a structural transition in the 48S from an open, scanning-competent conformation to a closed, scanning-arrested conformation occurs (*14*). This conformational change is accompanied by the release of eIF1, eIF2 and GDP, leaving the Met-tRNA_i_^Met^ at the P site of the 40S base paired with the AUG codon (*10*). A second GTP-regulated step, catalyzed by initiation factor eIF5B, is then required for the recruitment of the large (60S) ribosomal subunit (*15, 16*). A full (80S) ribosome primed with mRNA and Met-tRNA_i_^Met^ at the P site then transitions towards the fast, less regulated elongation phase (*17, 18*).

The above described pathway is referred to as the canonical, 5’-end and cap-dependent translation route of initiation (*12*). The bulk of eukaryotic mRNAs transitions this route, however, deviations from the canonical route are common, normally associated with translation under stress conditions (*19, 20*). Usually, non-canonical initiation is associated with extended <u>5’ UnTranslated Regions</u> (5’-UTRs) on mRNAs (*21, 22*). In complex eukaryotes, 5’-UTRs can be very long and can harbor short <u>O</u>pen <u>R</u>eading <u>F</u>rames (ORFs) designated as upstream ORFs or uORFs (*22, 23*). These uORFs are enigmatic, as ribosome-profiling experiments clearly show ribosome positioning on them, however, to date, the short peptides encoded in uORFs could have not been unambiguously identified by mass-spectrometry (*24*).

Well studied examples of the functional relevance of uORFs at 5’-UTRs in stress regulated genes, can be found in the yeast stress response regulator GCN4 or the mammalian transcription factor ATF4 (*25, 26*). In these stress regulated genes, the presence of several uAUG codons has been shown to be essential for differential translation regimes in homeostasis versus stress conditions (*22*). Other examples of translation regulation by uAUG codons are less understood, for example, uAUG codons immediately followed by a stop codon (designated as “start-stop uORFs”) are also found in 5’-UTRs of mammalian mRNAs (*23, 27*). Re-initiation events involving translation termination factors have been suggested to play a role in regulating start-stop uORFs, but in general, the function and possible biological activities of the peptides encoded in uORFs or the mechanism that mediate “start-stop uORFs” regulation remains obscure (*26, 28*).

A better characterized example of non-canonical initiation can be found in viral mRNAs (*29*). Viruses exploit the complexity of eukaryotic initiation to gain access to the host machinery for protein production (*30*). Strategies like mimicking the cap structure or transferring caps from cellular mRNAs (“cap-snatching”) allow viral mRNAs to hijack host ribosomes, redirecting them towards the production of viral proteins (*6, 30*). Another viral strategy for ribosome hijacking is the use of structured RNA sequences in viral mRNAs (*31*). These sequences are called Internal <u>R</u>ibosomal <u>E</u>ntry <u>S</u>ites (IRES) and a tentative classification based on their degree of RNA structure and dependency on canonical initiation factors, divided them in four main types (*32, 33*). Type I IRESs are low structured IRESs requiring all canonical initiation factors plus additional proteins not involved in canonical translation (called ITAF: IRES *<u>T</u>rans* <u>A</u>ctivator <u>F</u>actors) for efficient initiation. These IRESs also require scanning to locate the initiation codon. Type II IRESs present similar requirements of initiation factors as the Type I family, dispensing only with eIF1 and eIF1A as no scanning is required in this family. The assembly of initiation complexes is directly achieved on the AUG codon. Type III IRESs are more structured IRESs that fold in a compact structure able to interact with the 40S and the only required initiation factors are eIF2 and eIF3. Finally, the type IV family of viral IRESs is represented by highly structured IRESs with no requirement for canonical initiation factors to assemble translating ribosome on viral mRNAs.

The *Dicistroviridae* family of positive single-stranded RNA ((+)-ssRNA) viruses employs two types of IRESs to differentially express the regulatory versus the structural genes (*34*). The genome architecture of these viruses functionally segregates both kind of genes in two ORFs (Fig. 1A) (*35*). The first ORF is preceded by an approximately 700 nucleotides long 5’-UTR which harbors an IRES assigned to the type III family (*36*). *In vitro* characterization of the 5’-UTR-IRES of the <u>Cr</u>icket <u>P</u>aralysis <u>V</u>irus (CrPV), a prototypical *Dicistrovirus,* narrowed down the region of the 5’-UTR responsible for the IRES activity and established the strict requirement of eIF3 for this IRES to initiate translation. Interestingly, the AUG codon of the CrPV ORF1 is immediately preceded by a “start-stop uORF” (*36*). The combination of these two non-canonical initiation resources, namely a structured IRES and a “start-stop uORF” at the 5’-UTR, within the same viral mRNA, poses the question of how translation initiation is achieved by this RNA.

**Figure 1.**
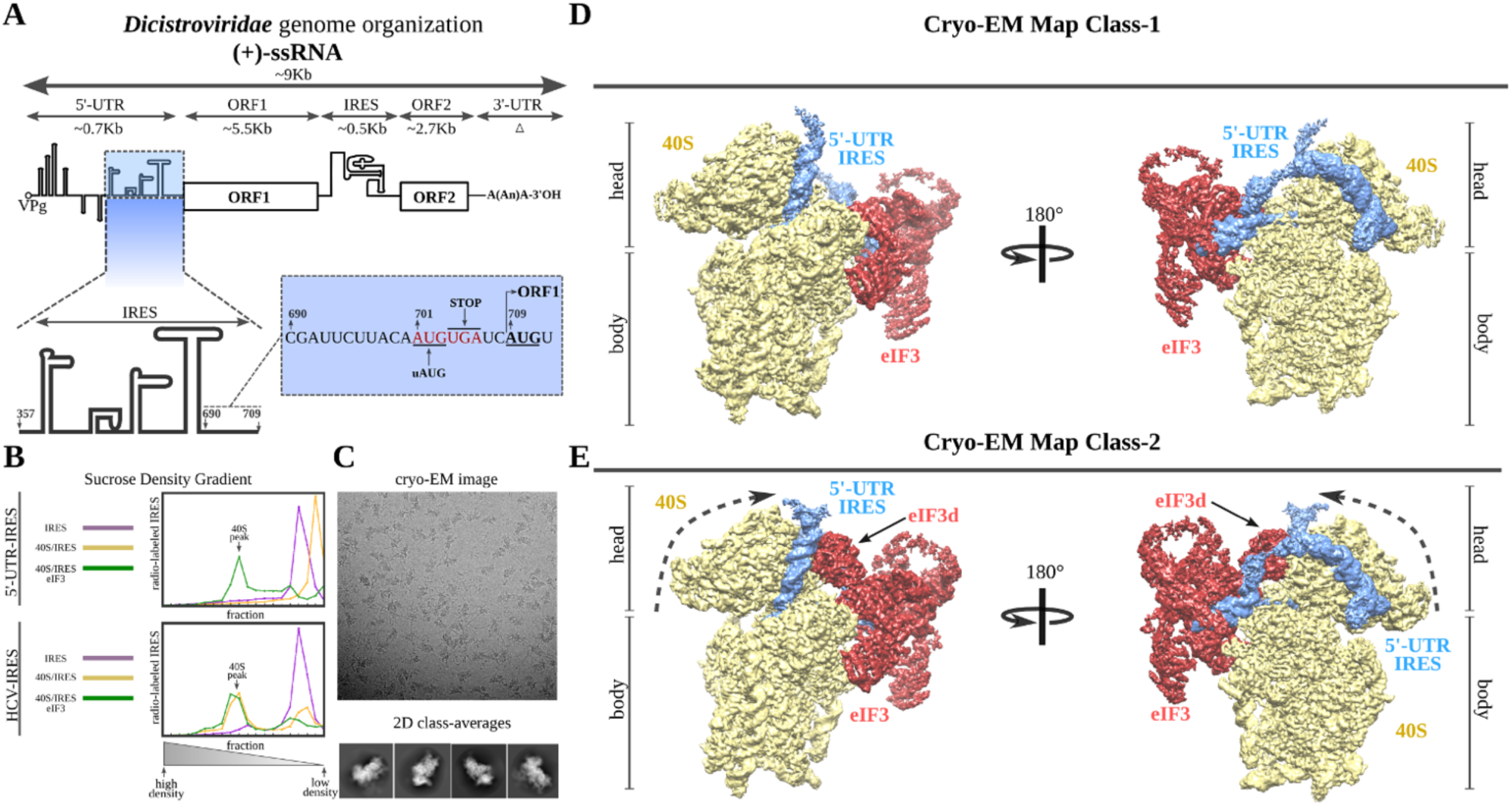
*Dicistroviridae* genome organization, *in vitro* complex formation and cryo-EM maps. **A,** Top, schematic representation of the genome organization of *Dicistroviruses*. Indicated with arrows are approximate genomic lengths for the different components. Bottom, detailed view of the region described to harbor the IRES activity of the 5’-UTR of the CrPV. On the right, sequence adjacent to the initiation AUG codon of ORF1, located at nucleotide 701 and preceded by a “start-stop uORF” indicated in red. **B**, Sucrose-gradient analysis showed the dependency on eIF3 of the 5’-UTR-IRES in order to form a stable complex with the 40S. Only in the presence of eIF3, 5’-UTR-IRES co-migrates with the 40S (top). In contrast, HCV IRES does not require eIF3 for 40S binding (bottom). **C**, Representative cryo-EM micrograph of the 40S/5’-UTR-IRES/eIF3 complex. Bottom, representative reference-free 2D class averages used for further image processing. **D**, **E** After 3D classifications, two classes showing density for 40S (yellow), eIF3 (red) and 5’-UTR-IRES (blue) could be found in the data set. Class-1 (top, **D**) presents a non-swiveled configuration of the 40S head and density for eIF3d is absent. Class-2 (bottom, **E**) shows a swiveled configuration of the 40S head (arrows) with eIF3d (indicated) contacting eIF3’s core subunits.

We sought to structurally characterize the 5’-UTR-IRES of the CrPV in its ribosome bound configuration, to gain insights on the ribosome binding determinants of this peculiar IRES as well as to understand how the delivery of Met-tRNA_i_^Met^ is accomplished. Two high resolution cryo-EM reconstructions of 40S/5’-UTR-IRES/eIF3 complexes combined with biochemical analysis, allowed us to precisely characterize how this IRES, using an extended structure with a modular, multi-domain architecture, binds and manipulates the 40S.

These results stress the importance of the unstudied diversity of viral IRESs and expand our understanding on the role 5’-UTR regions play in eukaryotic translation.

## Results

### The 5’-UTR-IRES of the CrPV requires eIF3 for a stable interaction with the 40S

Previous studies of the IRES located at the 5’-UTR of the CrPV (5’-UTR-IRES from hereafter) precisely defined the region of the 5’-UTR responsible for the IRES activity (residues 357 to 709) as well as its dependency on initiation factor eIF3 for efficient translation initiation (*36*). Sequence alignments of 5’-UTR regions of different viruses from this clade as well as with other viruses of the same family failed to identify any similarity with described IRESs of the type III family, like the <u>H</u>epatitis <u>C V</u>irus IRES (HCV-IRES) or the <u>C</u>lassical <u>S</u>wine <u>F</u>ever <u>V</u>irus IRES (CSFV-IRES) (*37, 38*). In contrast with the well characterized type IV family of IRESs found in the <u>I</u>nter<u>G</u>enic <u>R</u>egion (IGR-IRES) of these viruses, where strong sequence conservation allows comparisons and identification of possible structural motifs, the 5’-UTRs of *Dicistroviruses* seem to harbor divergent sequences, making structural modelling based on sequence conservation difficult (*39*). In order to address this gap in knowledge, we produced a truncated version of the 5’-UTR region of the genomic RNA of the CrPV containing the IRES (residues 357 to 728, Fig. 1A) to obtain structural information of its 40S bound conformation by electron cryo-microscopy (cryo-EM). We initially tested the *in vitro* dependency of 5’-UTR-IRES on eIF3 to engage purified 40S ribosomal subunits in a stable interaction. We assayed the ability of the 5’-UTR-IRES to co-migrate with purified 40S in sucrose density gradients as a test for the presence of a stable complex suitable for structural studies (Fig. 1B). Unexpectedly, the 5’-UTR-IRES does not form a stable complex with the 40S in the absence of eIF3, in contrast to the HCV-IRES, which is able to form stable complexes with the 40S subunit alone and even with full (80S) ribosomes (Fig. 1B) (*37*). In the presence of eIF3, however, the 5’-UTR-IRES co-migrates with purified 40S subunits, demonstrating the presence of a stable complex (Fig. 1B). This complex revealed clear particles in cryo-EM images, rendering detailed two-dimensional class averages where density for eIF3 could be identified albeit at lower threshold (Fig. 1C). The 40S/5’-UTR-IRES/eIF3 complex exhibited a delicate behavior under cryo-EM conditions, with a strong tendency to disassemble in thin ice. Extensive screening for suitable ice areas was essential to obtain particles of the fully assembled complex (Fig. S1A and B). The sample also exhibited a high degree of heterogeneity, that could be resolved by image processing in Relion (*40, 41*) (Fig. 1D,E and Fig. S1C).

Two main classes of particles containing density for 5’-UTR-IRES, 40S and eIF3 were found in the dataset (Fig. 1D and E). Both classes contain density for the 40S, the IRES and the core subunits of eIF3 (a/c/e/k/l/f/m) and class-2 additionally, presents density for eIF3 subunit d (Fig. 1E, eIF3d). Class-2 also exhibits a 40S head in a swiveled configuration. The eIF3d subunit follows the 40S head swiveling movement to establish interactions with eIF3a, a core subunit of eIF3 (see below).

Robust density ascribable to the 5’-UTR-IRES could be found in both classes (Fig. 1D and E, blue). The ribosome bound conformation of 5’-UTR-IRES shows an extended configuration, circling the 40S head (Fig. 2A). Three domains connected by flexible linkers, could be defined: an elongated domain I (DI) at the back of the 40S head contacting ribosomal proteins uS3 and RACK1 (Fig. 2), a second domain (DII) formed by a dual hairpin at the back of the 40S body interacting with eIF3 (Fig. 3) and a third, large helical domain (DIII) placed at the periphery of the 40S E site, contacting ribosomal proteins uS7 and uS11 (Fig. 4).

**Figure 2.**
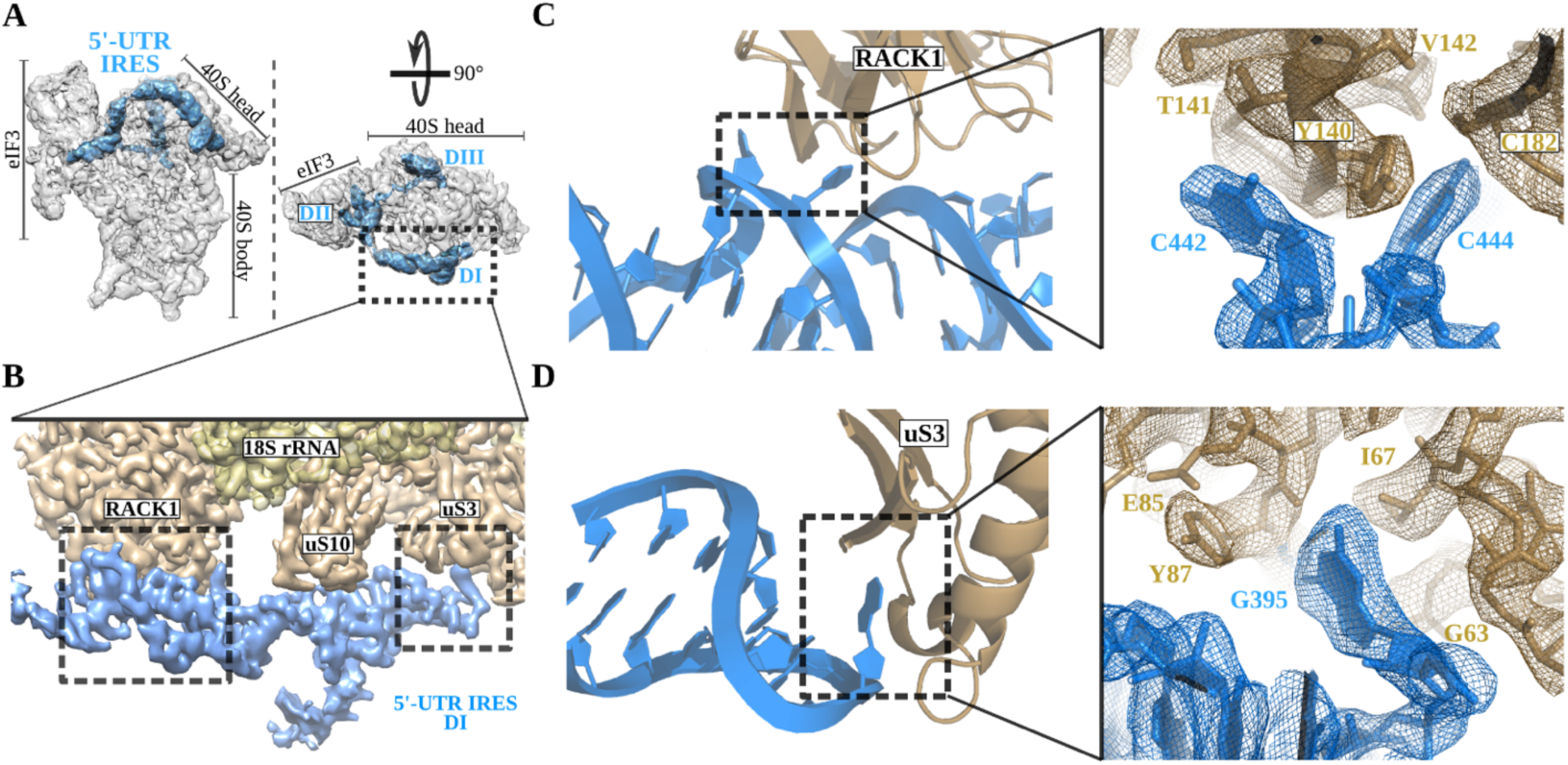
5’-UTR-IRES domain I engages ribosomal proteins RACK1 and uS3. **A,** Overview of the 40S/5’-UTR-IRES/eIF3 map with 40S and eIF3 depicted grey and 5’-UTR-IRES blue. **B**, Detailed view of the cryo-EM density of the 40S/5’-UTR-IRES/eIF3 map centered around 5’-UTR-IRES domain I (DI). Ribosomal proteins are colored dark yellow, 18S rRNA yellow and 5’-UTR-IRES blue. Contacts between 5’-UTR-IRES domain I and ribosomal proteins RACK1 (**C**) and uS3 (**D**) could be defined due to excellent local cryo-EM densities.

**Figure 3.**
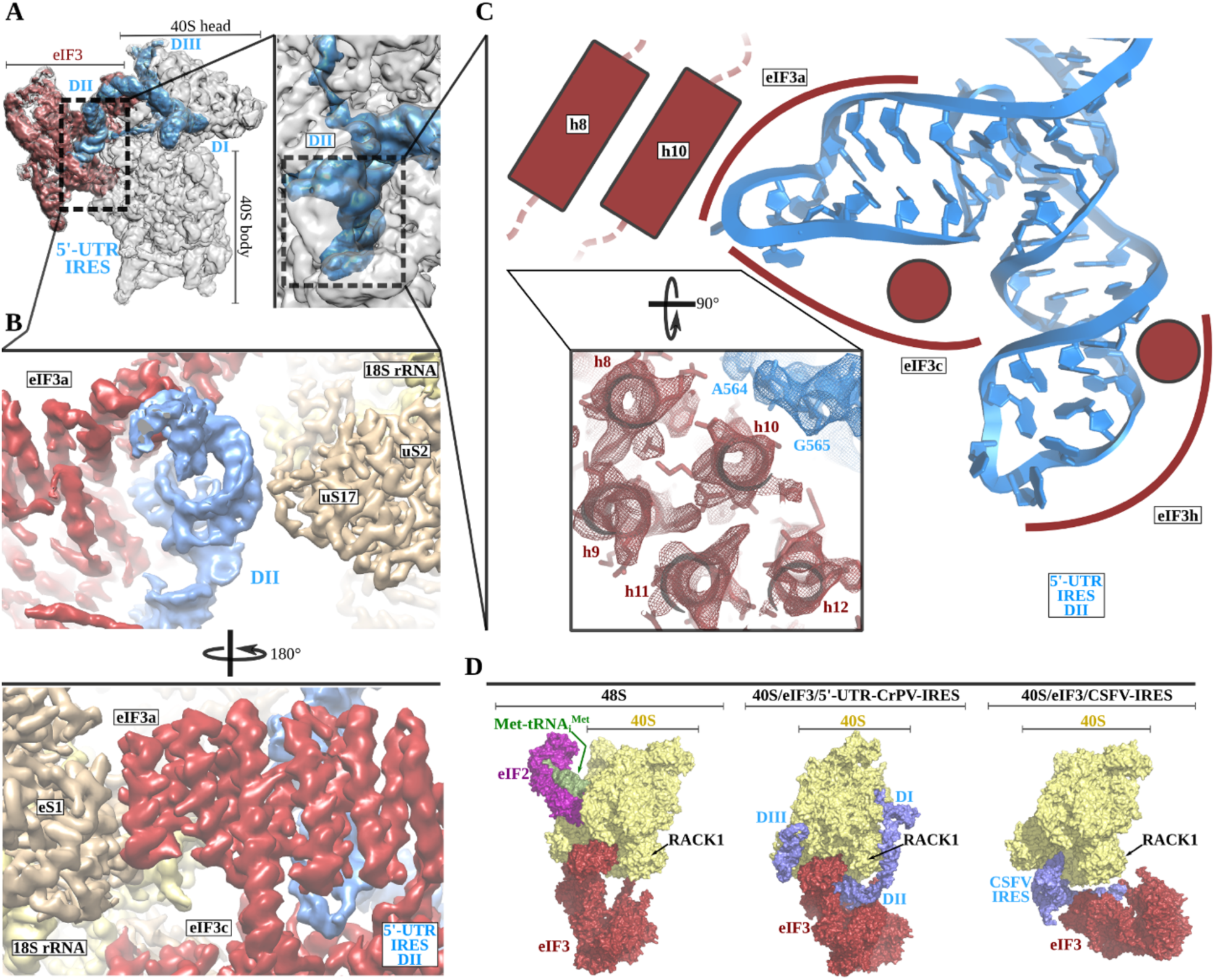
5’-UTR-IRES domain II is formed by a dual hairpin mediating eIF3 recruitment. **A,** Overview of the 40S/5’-UTR-IRES/eIF3 cryo-EM map with 40S colored grey, eIF3 red and 5’-UTR-IRES blue. On the right a zoomed view centered around 5’-UTR-IRES domain II. **B**, Detailed view of the cryo-EM map for the region occupied by 5’-UTR-IRES domain II with 40S components colored yellow, eIF3 red and 5’-UTR-IRES blue. Domain II is sandwiched between ribosomal protein uS17 located at the back of the 40S body and eIF3 core subunits a and c. **C**, 5’-UTR-IRES domain II is formed by a dual hairpin that establishes interactions with α-helices 8 and 10 from eIF3a. These contacts are mediated mainly by basic residues of eIF3 and the phosphate backbone of the IRES. **D**, superposition of the 40S/5’-UTR-IRES/eIF3 complex with canonical 48S complex (left, PDB ID 6FEC) and with the CSFV-IRES/40S complex (right, PDB ID 4c4q). The 5’-UTR-IRES binds to the 40S with a conformation compatible with the canonical position described for eIF3 in the 48S complex.

**Figure 4.**
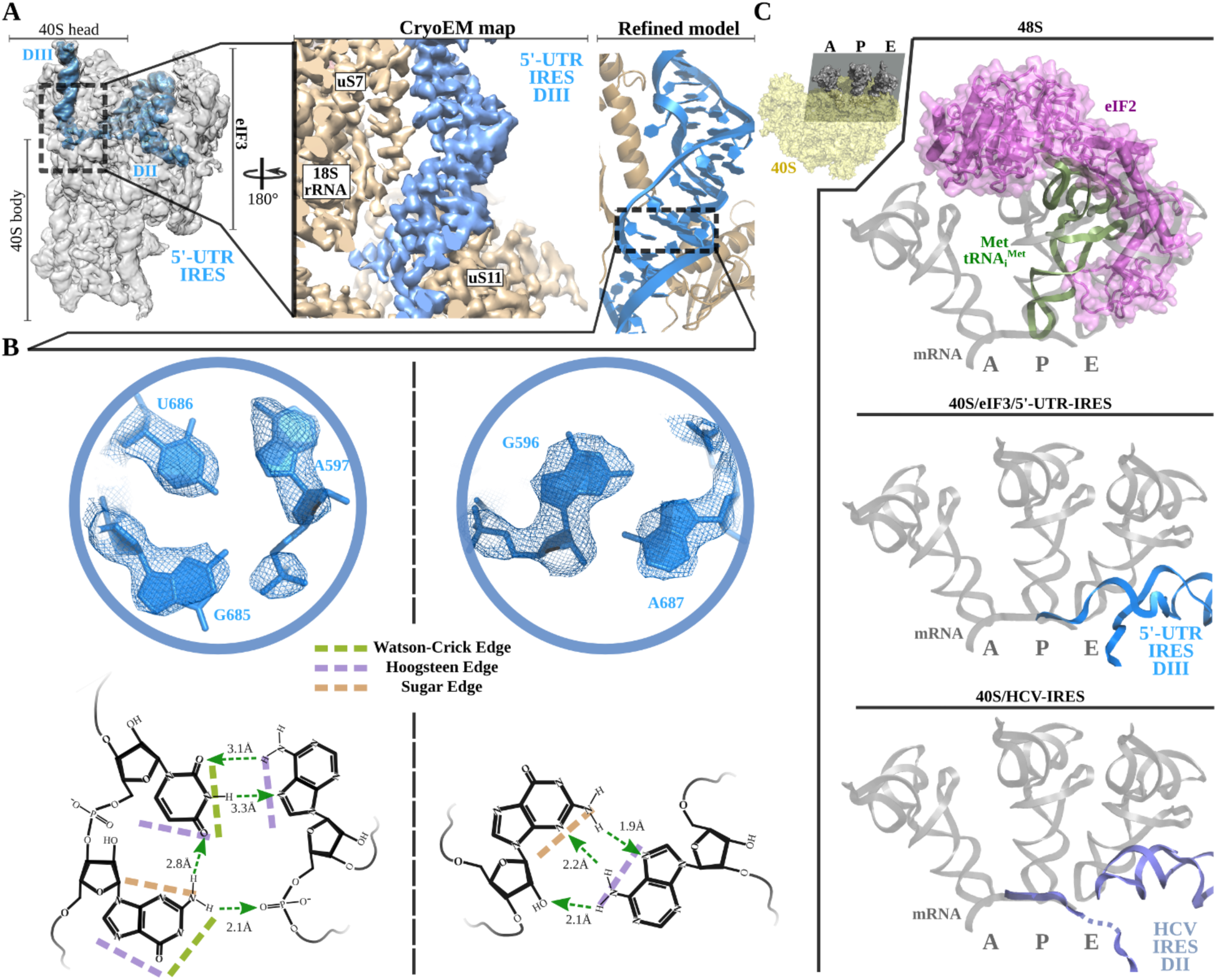
Non-canonical base pairing in 5’-UTR-IRES domain III assists on P site access. **A,** Overview of the 40S/5’-UTR-IRES/eIF3 cryo-EM map with 40S and eIF3 colored grey and 5’-UTR-IRES blue. On the right, detailed view of the E site where 5’-UTR-IRES domain III is placed, shown is the cryo-EM map with the 5’-UTR-IRES colored blue and 40S components yellow and, on the right, the final refined model colored following the same color scheme. Ribosomal proteins uS7 and uS11 as well as several 18S rRNA bases contact 5’-UTR-IRES domain III. **B**, Non-canonical base pairs found in 5’-UTR-IRES domain III induce a distortion of the double helix near the E site. Two examples are shown with the refined model inserted in the experimental cryo-EM density at the top, and corresponding chemical diagrams below with the base edges and hydrogen bonds involved in interactions indicated. **C**, Superpositions of canonical 48S complex (PDB ID 6FEC, top), HCV-IRES/40S complex (PDB ID 5A2Q, bottom) with the 40S/5’-UTR-IRES/eIF3 (middle) model focused on the tRNA A, P and E binding sites. 5’-UTR-IRES domain III and HCV-IRES occupies a space on the E site that overlaps with the position described for eIF2 in the canonical 48S complex.

### Domain I of the 5’-UTR-IRES contacts ribosomal proteins RACK1 and uS3

The 5’ proximal segment of the 5’-UTR-IRES (residues 357 to 486) forms the domain I, characterized by an elongated T-shaped structure anchored to the back of the 40S head (Fig. 2A and B). A long helical segment in this domain “wraps” around the apical part of ribosomal protein RACK1. Two bases of this helical segment of domain I, C442 and C444, are extruded from the body of the double helix to establish hydrophobic interactions with tyrosine residue 140 of RACK1 (Fig. 2C). These interactions bend the main helical segment of the 5’-UTR-IRES DI directing the tip of this domains towards ribosomal protein uS3 (Fig. 2D). Guanine residue 395 is inserted deep in a hydrophobic pocket of ribosomal protein uS3, establishing contacts with main chain atoms of this ribosomal protein. In this location, 5’-UTR-IRES DI is found adjacent to the mRNA entry channel of the 40S, overlapping with the space previously described to be occupied by the helicase DHX29 involved in canonical initiation (Fig. 2D) (*42*).

### 5’-UTR-IRES binding to the 40S is compatible with a canonical configuration of eIF3

The second domain of 5’-UTR-IRES (DII) is connected to domain I by a flexible linker poorly defined in our maps as it is exposed to the solvent. This second domain of the 5’-UTR-IRES is formed by a dual hairpin and is wedged between the back of the body of the 40S and eIF3 subunits a and c (Fig. 3A and B). Ribosomal protein uS17 peripherally contacts this domain, establishing interactions with the phosphate backbone of the IRES (Fig. 3B). A network of interactions involving eIF3 subunits a, c and h anchors DII to this position (Fig. 3C). These interactions are also established through contacts between negatively charged residues on eIF3 and the phosphate backbone of the IRES. No contacts with specific IRES bases could be observed.

Currently, medium resolution cryo-EM reconstructions for 40S complexes containing eIF3 and the rest of components of the canonical 48S complex are available (*43*) (PDB ID 6FEC) as well as for the 40S in complex with eIF3 and the CSFV-IRES (*38*)(PDB ID 4c4q). Comparisons of these structures with our complex, reveals a positioning of eIF3 relative to the 40S very similar to the canonical 48S complex and different from the position adopted by eIF3 in the CSFV-IRES/40S complex (Fig. 3D). In the 48S canonical configuration, eIF3 contacts the 40S through helix 1 of eIF3a and helix 22 of eIF3c as well as eIF3d which is isolated in its 40S interaction, away from the core subunits of eIF3 (Fig. 3D, left). The CSFV-IRES engages the 40S displacing eIF3 from its position in the canonical 48S. (Fig. 3D, right). Additionally, in the canonical 48S complex, eIF3 interacts with the 40S peripherally, allowing the presence of cavities between eIF3 and the back of the 40S. These cavities are exploited by the 5’-UTR-IRES which inserts its domain II in one of these cavities, adopting a configuration compatible with the binding of eIF3 to the 40S in the canonical 48S complex (Fig. 3D, middle). No major rearrangement of eIF3 (compared to its position in the canonical 48S complex) are required for the binding of the 5’-UTR-IRES, what could represent an advantage in hijacking preformed cellular 48S complexes ready to transit the cap-dependent route of initiation.

### Non-canonical base pairing in the 5’-UTR-IRES DIII places the uAUG codon near the P site

Threading through the 40S channel formed by ribosomal proteins uS7 and uS11, a flexible single stranded linker connects DII with DIII (Fig. 4A). DIII forms a prominent, helical mass in the surroundings of the E site of the small subunit at the inter-subunit face of the 40S. The helical segment is very well defined in our maps as it is stabilized by numerous contacts with ribosomal proteins uS7, uS11 and 18S ribosomal RNA (rRNA) bases (Fig. 4 **and** Fig. S3). However, the distal part of this domain forms two short stem loops that given their flexibility could only be modelled at low resolution.

Inspection of the cryo-EM density in both classes reveled a distortion in the canonical double helix of the main segment of this domain as it approaches the E site. The quality of the maps in this area allowed *de novo* modelling of these residues, revealing a set of non-canonical interactions between the RNA bases (Fig. 4A and B). In-plane triple base interactions involving sugar and Hoogsteen edges of the bases as well as purine-purine Hoogsteen base pairs could be found in this stretch of residues of the helical segment of DIII (Fig. 4B)(*44*). Overall, these non-canonical base pairs induce a distortion at the base of DIII helping in the positioning of the single stranded segment of the 5’-UTR-IRES harboring the uAUG codon at position 701, close to the P site (Fig. 4C, middle). The 5’-UTR-IRES thus accesses the 40S P site through the E site, what precludes, in the present configuration, a concurrent recruitment of the TC (eIF2/Met-tRNA_i_^Met^/GTP, Fig. 4C). Interestingly, a similar strategy is followed by the HCV-IRES; a superposition of the structure of the HCV-IRES in complex with the 40S (*45, 46*)(PDB ID 5A2Q) with our structure reveals a very similar positioning of the domain II of HCV-IRES, accessing the P site through the E site to position the AUG codon in the surroundings of the P site (Fig. 4C, bottom). Even though both IRESs differ markedly in their interaction with the back of the 40S and eIF3, both converge to similar structural solutions for the placement of the AUG initial codon close to the 40S P site. This fact could be rationalized as the A/P/E ribosomal sites are highly conserved in all kingdoms of life whereas ribosomal proteins, especially eukaryotic specific ones, used for the docking of these IRESs to the back of the 40S, present less degree of conservation and it would not be unexpected different IRESs explore alternative solutions for their binding.

### Swiveling of the 40S head locks the 5’-UTR-IRES inducing a compact conformation of eIF3

Initial processing of the cryo-EM data revealed a marked dynamic of the head of the 40S. Masked classification and refinement in Relion3 (*47*) revealed two mayor populations, distinguishable by different degrees of swiveling of the 40S head (Fig. 1 and Fig. S1C). The 40S head is attached to the body by a single RNA helix, making this component of the ribosome extremely flexible (*33*). Intrinsic and independent movements of the 40S head are instrumental in tRNA translocation and also in canonical initiation (*48, 49*). The 5’-UTR-IRES seems to exploit this intrinsic dynamic to, in a first instance, bind to the 40S and, in a second instance, “lock” the IRES in a specific conformation committing the complex towards viral translation (Fig. 5). In the class-1 (open conformation) the head of the 40S shows almost a canonical configuration with very little swiveling and no tilt. In this conformation, the latch of the 40S (an early defined contact between the head and the body of the 40S (*50*)) is closed. At the other side of the 40S head, access to the channel formed by ribosomal proteins uS7 and uS11 is exposed and eIF3d density is not well defined, probably due to a high degree of flexibility or low occupancy (Fig. 5A, left). In class-2 (closed conformation), the head of the 40S has experienced a medium range degree of swiveling, compared to the widest displacement reported (*48*). The movement of the 40S head is followed by the 5’-UTR-IRES, with DI experimenting the highest displacement compared to its position in class-1 (Fig. 5A, right and B). In the swiveled conformation, the latch is open, and the channel formed by ribosomal proteins uS7 and uS11 is plugged by eIF3d, which in this class, presents robust density (Fig. 5C). These movements are restricted to the 40S-head and the domains of 5’-UTR-IRES, especially DI. No movement could be detected in the core subunits of eIF3, maintaining in the closed class an identical configuration respect the 40S body as in the open class (Fig. 5B). The swiveled configuration of the 40S brings eIF3d close to eIF3a, one of the core subunits of eIF3 (Fig. 5C). Well defined density in this area could be observed for the interface eIF3a/eIF3d (Fig. 5C, right). Thus, in the closed conformation, eIF3 shows a hitherto unknown conformation, which we speculate could reflect a transient state populated at some point during canonical initiation and exploited by the 5’-UTR-IRES in order to gain access to the P site of the small subunit (*51*).

**Figure 5.**
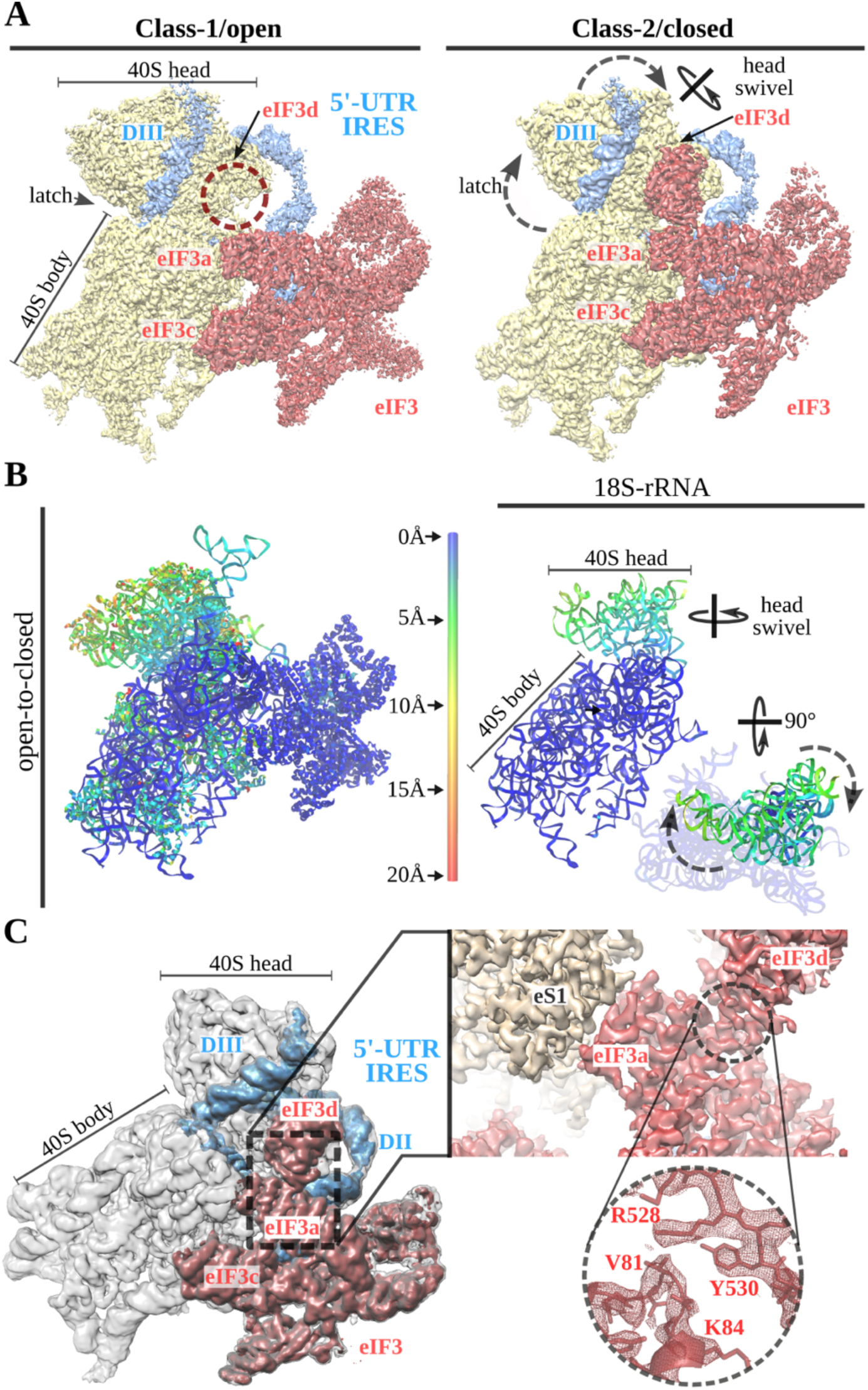
A 40S head swiveling movement “locks” 5’-UTR-IRES on the 40S inducing a compact eIF3 configuration. **A,** Cryo-EM maps obtained for the two classes present in the 40S/5’-UTR-IRES/eIF3 dataset with 40S colored yellow, eIF3 red and 5’-UTR-IRES blue. Indicated, the position of the latch, eIF3d and the swiveling rotation axis. A 40S head swiveling movement in class-2 brings closer eIF3d to the core subunit of eIF3, eIF3a, establishing interactions that stabilize its conformation. **B**, Left, ribbon diagram colored according to pairwise root <u>m</u>ean <u>s</u>quare <u>d</u>eviation (r.m.s.d.) displacements for the open to close transition with displacements scale at the center. On the right, simplified diagram showing only the 18S rRNA colored with the same scale as on the left. Two orthogonal view are shown where it can be appreciated the main displacement is localized on the 40S head. **C**, Overview of the closed class with 40S colored grey, eIF3 red and 5’-UTR-IRES blue. Inset, detail of the experimental density obtained for the eIF3a/eIF3d interface for this class. Clear side chains information was present in the maps, allowing proper building and model refinement.

### TC delivers Met-tRNA_i_^Met^ to uAUG at position 701 and initiation factors eIF1 and eIF1A assist on AUG location

Our structures of the 40S/5’-UTR-IRES/eIF3 complex reveled a positioning of the DIII of the IRES that overlaps with the position the TC populates at the E site in canonical initiation (Fig. 4C) (*14, 43*). Additionally, in our maps, we could only confidently identify density for the single stranded segment of RNA of the IRES placed close to the P site until residue 695, whereas the canonical AUG of ORF1 is found at nucleotide 709. These facts prompted us to wonder how the delivery of Met-tRNA_i_^Met^ to the AUG is accomplished. Making use of reconstituted initiation assays with native components and toe-printing analysis (*52*), we could dissect the different steps followed by the 5’-UTR-IRES in order to correctly place Met-tRNA_i_^Met^ based paired with the AUG codon (Fig. 6A). TC by itself is able to load Met-tRNA_i_^Met^ on the 40S/5’-UTR-IRES/eIF3 complex in isolation (Fig. 6A, lane 2). Interestingly, this loading event is not directed towards the canonical AUG but to the uAUG located at position 701 which is a part of the “star-stop uORF” that precedes the *bona fide* AUG codon of ORF1 (Fig. 1A). A similar uAUG delivery of Met-tRNA_i_^Met^ can be accomplished by eIF5B which, under stress conditions, has been described to substitute eIF2 for Met-tRNA_i_^Met^ delivery (*53–55*), following then eukaryotic initiation a “bacterial-like” mode of initiation (Fig. 6A, lane 4). Transitioning to the correct AUG could only be detected in the presence of eIF1/eIF1A but only when the TC was present and not for eIF5B (Fig. 6A, lanes 3 and 5). Notably, the presence of eIF1/eIF1A seems to be detrimental for uAUG Met-tRNA_i_^Met^ loading by eIF5B as their presence significantly reduces the toe-print signal that can be observed for eIF5B in isolation. However, no concomitant increase in toe-print signal for the canonical AUG could be observed for the eIF1/eIF1A/eIF5B reaction. Only eIF2 as part of the TC and assisted by eIF1/eIF1A can properly locate the *bona fide* AUG of ORF1, probably through an early 48S intermediate assembled at the uAUG of the “star-stop uORF” located at nucleotide 701.

**Figure 6.**
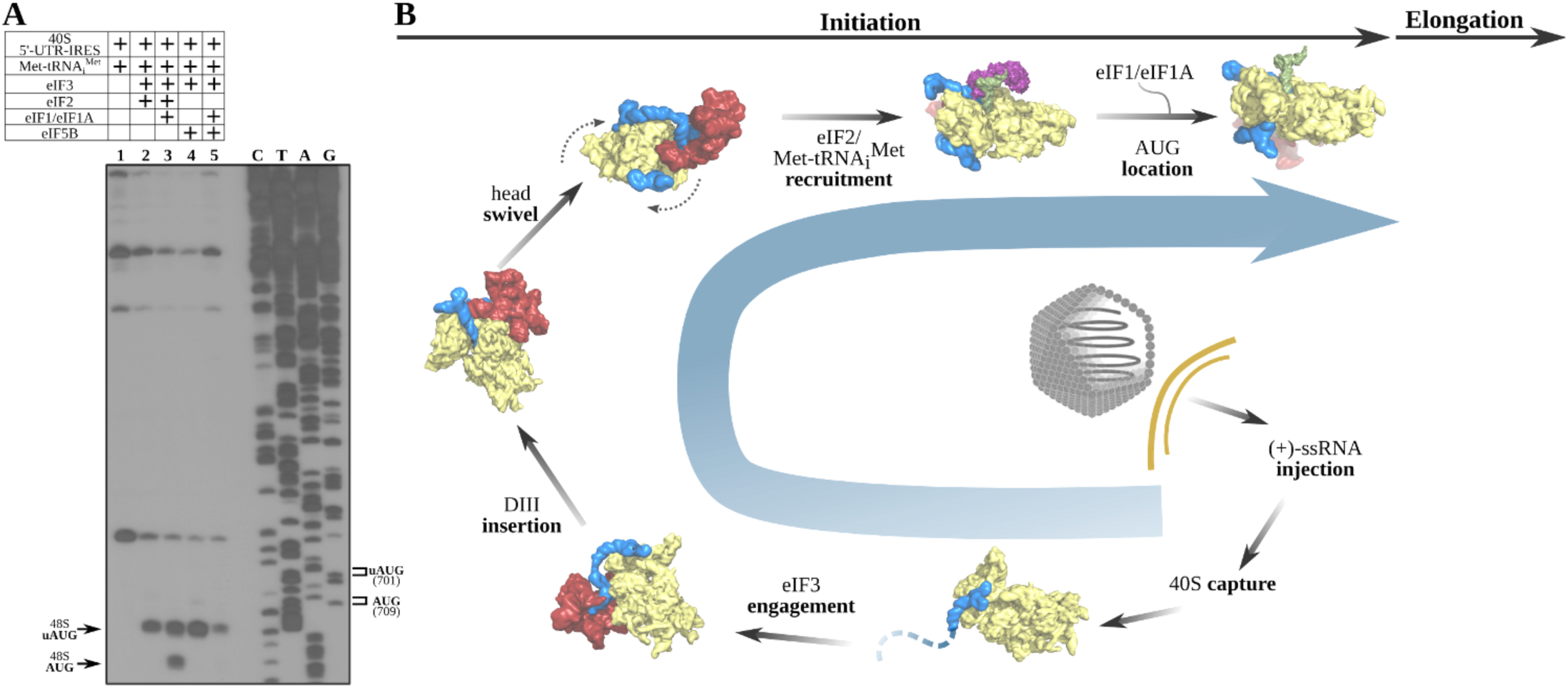
5’-UTR-IRES requires TC, eIF1 and eIF1A to assemble a functional initiation complex via an uAUG intermediate. **A,** Toe-print analysis of 48S initiation complexes assembled on 5’-UTR-IRES in an *in vitro* reconstituted system. eIF2 delivers Met-tRNA_i_^Met^ to the uAUG (lane 2) and requires the presence of eIF1/eIF1A to transition to the *bona fide* AUG codon of ORF1 (lane 3). eIF5B, under some conditions, can substitute eIF2 in Met-tRNA_i_^Met^ delivery (*53*). In the absence of eIF1/eIF1A, a robust toe-print signal is detected in the presence of eIF5B (lane 4), however, Met-tRNA_i_^Met^ is delivered to the uAUG, thus eIF5B is unable to find the annotated AUG even in the presence of eIF1/eIF1A (lane 5). **B**, A model for 5’-UTR-IRES mediated translation initiation, from the bottom right: injection of the genomic (+)-ssRNA of the CrPV into the cytosol allows the pre-folded 5’-UTR-IRES DI to capture free 40S subunits. Via interactions with DII, eIF3 is recruited to the complex with an initial canonical conformation of the 40S head without tilt or swivel. Insertion of 5’-UTR-IRES DIII in the vicinity of the E site and a swiveling movement of the 40S head, induce a “locked” conformation of the complex with the uAUG at 701 in the vicinity of the P-site. Delivery of Met-tRNA_i_^Met^ and location of ORF1 AUG is achieved by the concerted action of eIF2, eIF1 and eIF1A via an uAUG intermediate. Large subunit recruitment mediated by eIF5B will allow transitioning into elongation.

## Discussion

The development of ribosome profiling techniques and its application to eukaryotic samples, specially mammals, have expanded our once simplified perception of translational landscapes (*24, 56*). There is now direct evidence that translation regulation can be accomplished following many different and alternative pathways, involving not only direct regulation of ribosomes but also adjacent sequences to the coding regions of mRNAs (*57*). 5’-UTRs are of special interest, given their pervasive presence in mammalian transcriptomes (*21, 58*). Ribosome-profiling datasets have revealed the presence of translating ribosomes paused on 5’-UTRs, implying a decisive role in regulating translation, specially under stress conditions (*59, 60*). Given the proficiency viruses exhibit in exploiting the eukaryotic machinery for protein production, it would be surprising the plethora of regulatory mechanisms implemented at the level of 5’-UTRs would not be leveraged by viruses to hijack the eukaryotic apparatus for protein synthesis.

The 5’-UTR of the (+)-ssRNA of the *Dicistovirus* CrPV harbors an IRES able to direct initiation towards ORF1 in the early phase of infection (*35, 61*). Expression of ORF1 is instrumental for virus replication, as the RNA-dependent RNA polymerase (RdRp) and the protease responsible for the proteolytic digestion of the polyprotein containing the structural proteins are encoded in ORF1 (*62*). Given the poor sequence conservation between 5’-UTR regions within this family of viruses, it was challenging even to propose a secondary structure diagram for this IRES. Applying cryo-EM techniques combined with toe-print analysis of initiation reactions assembled with native components, we were able to visualize the three-dimensional structure of the 5’-UTR-IRES in its ribosome bound configuration and to describe the initiation route followed by this IRES to assemble a functional initiation complex competent for elongation.

The 5’-UTR-IRES features a novel multi-domain, extended architecture that encircles three quarters of the 40S head, exploiting binding sites not previously described for any IRESs (Fig. 2, 3 and 4). Ribosomal proteins uS3 and RACK1 are used by the IRES to anchor its DI to the back of the 40S head (Fig. 2). The structure thus rationalizes previous data showing a preeminent role of RACK1 in CrPV and related viruses infecting *Drosophila* (*63*). The interaction of DI with RACK1 is also instrumental to position DII at the back of the 40S body, sandwiched in between ribosomal protein uS17 and eIF3 (Fig. 3). Interestingly and in contrast with the HCV-IRES, the conformation observed for eIF3 in the complex with 5’-UTR-IRES is very similar to the conformation observed for eIF3 in the 48S complex, with the IRES “filling up” cavities present between the 40S and eIF3 in this canonical complex (*43*). The HCV-IRES and related IRESs like the CSFV-IRES displace eIF3 from its canonical location, using a very different mechanism for IRES docking to the 40S (*38*).

In order to place the AUG of ORF1 in the surroundings of the P site, the 5’-UTR-IRES accesses the P site through the E site, in a similar manner as the HCV-IRES (Fig. 4C) (*45*). In this aspect, the 5’-UTR-IRES recapitulates binding strategies known for other IRESs like the IGR-CrPV-IRES that also make use of ribosomal protein uS7 for its binding to the ribosome or the mentioned HCV-IRES which places its domains II and IV in the surroundings of the P site, sliding the elongated DII from the back of the 40S to the P site through the E site (*64*).

The placement of the AUG of ORF1 in the surroundings of P site seems to be exerted by a mechanism involving the intrinsic dynamics of the 40S head (*33*)(Fig. 5). The 5’-UTR-IRES exploits the characteristic swiveling movement of the 40S head to bind and progress towards a conformation that “locks” the IRES on the 40S and at the same time, induces a compact conformation of eIF3 with subunit eIF3d in close contact with the core subunits of eIF3. We speculate this hitherto unknown conformation of eIF3 may also be relevant for canonical initiation, as the cap binding protein eIF4E and the rest of initiation factors of the eIF4 family interact with eIF3 in the same region (*51*).

We propose the following comprehensive model for how the 5’-UTR-IRES of the CrPV operates: immediately after the (+)-ssRNA genomic molecule of the CrPV is injected in the cytoplasm of the host cell, the IRES harbored at the 5’-UTR captures 40S subunits by its DI (Fig. 6B, bottom right). Recruitment of eIF3 is mediated by DII, allowing the sliding of the flexible linker connecting DII and DIII between the head and the platform of the 40S to place DIII in the surroundings of the E site (Fig. 6B, bottom left). A swivel movement of the 40S head closes the channel between the head and the platform of the 40S effectively “locking” the 5’-UTR-IRES in the 40S, inducing a compact conformation of eIF3 with eIF3d subunit in interacting distance with eIF3’s core subunit a (Fig. 6B, left top). With this configuration, eIF2 as part of the TC can deliver Met-tRNA_i_^Met^ to the uAUG located at nucleotide 701 and further assistance by initiation factors eIF1/eIF1A allows for a downstream location of the AUG codon of ORF1 at nucleotide 709. Large subunit recruitment grants transitioning towards elongation, committing the ribosome to the production of viral proteins (Fig. 6B, right top).

The extent and diversity of viral populations contrast with our limited understanding of viral strategies for translation competency (*3*). Little is known regarding the structural diversity of viral IRESs or how 5’-UTR regions and accessory mRNA sequences are leveraged by viruses for ribosome hijacking (*65*). IRES studies have been limited to a handful of model viral IRESs because they offer advantageous traits, like compact conformations and structural stability (*31*). A preliminary classification of viral IRES sequences was proposed, dividing the known IRESs in four main families (*32*). However useful, this simplified classification may be limited in describing the vast diversity of IRES mechanisms found in nature. For example, the 5’-UTR-IRES of the CrPV described here was assigned to type III family of IRESs given its dependency on eIF3 and eIF2. This family of IRES is also defined by the direct assembly of initiation complexes in AUG initiation codons, without the need for “scanning” (*12*). However, the 5’-UTR-IRES assembles an initiation complex via an intermediate at a uAUG codon, and then, transition to the *bona fide* AUG codon with the assistance of eIF1 and eIF1A. Even though we cannot call this transition a proper “scanning” event, it isn’t either a direct assembly on the AUG codon, what challenges the definition for type III IRESs.

In summary, we have structurally characterized the 5’-UTR-IRES of the CrPV in its ribosome bound state and discovered a novel mechanism of translation initiation employing an early intermediate assembled on a uAUG codon. Given the rich diversity of viral sequences in the animal virome, new IRESs exploiting different aspects of animal translation will be probably discovered.

## Acknowledgements.

We are grateful to Dr. Jean-Luc Imler for a generous donation of a CrPV-5’-UTR-IRES plasmid. We are also thankful to Prof. Jennifer Doudna for a generous donation of a HCV-IRES transcription vector. We are thankful to Bob Grassucci and Zhening Zhang for assistance in cryo-EM data acquisition. Part of this work was performed at the Simons Electron Microscopy Center and National Resource for Automated Molecular Microscopy located at the New York Structural Biology Center, supported by grants from the Simons Foundation (SF349247), NYSTAR, and the NIH National Institute of General Medical Sciences (GM103310).

## Methods

### 5’-UTR-IRES and HCV IRES production

For cryo-EM analysis, a transcription vector for 5’-UTR-IRES (nucleotides 357-728) was constructed inserting a T7 promoter sequence upstream of 5’-UTR-IRES sequence followed by an BamHI restriction site, using pUC19 as a scaffold vector. For toe-print assay, 5’-UTR-IRES with the extended ORF part for primer annealing was cloned employing a similar strategy. T7 RNA polymerase *in vitro* transcription and purification on Spin-50 mini-column (USA Scientific) were used to obtain highly purified 5’-UTR-IRES and HCV-IRES RNAs.

### Purification of translation components and ribosomal subunits

Native 40S subunits, eIF2, eIF3, eIF5B and rabbit aminoacyl-tRNA synthetases were prepared as previously described (*66*). Recombinant eIF1 and eIF1A were purified according to a previously described protocol (*52*)*. In vitro* transcribed Met-tRNA_i_^Met^ was aminoacylated with methionine in the presence of rabbit aminoacyl-tRNA synthetases as previously described (*67*).

### Assembly of ribosomal complexes

To reconstitute different ribosomal complexes for toe-print assay, we incubated 0.3 pmol 5’-UTR-IRES RNA with 1.8 pmol 40S subunits, 10 pmol eIF1, 10 pmol eIF1A, 10 pmol eIF2, 5 pmol eIF3, 5 pmol eIF5B, and 5 pmol Met-tRNA_i_^Met^, as indicated, in a 20 uL reaction mixture containing buffer A (20 mM Tris-HCl, pH 7.5, 100 mM KCl, 2.5 mM MgCl_2_ and 1 mM DTT) with 0.4 mM GTP and 0.4 mM ATP for 10 min at 37◦C. We analyzed the assembled ribosomal complexes via a toe-print assay essentially as described (*66*). For sucrose density gradient experiment, we incubated co-transcriptionally [32-P]-labelled 5’-UTR-IRES or HCV-IRES RNAs with 3.7 pmol 40S subunits and 11 pmol eIF3, as indicated, in a 60 uL reaction mixture containing buffer A for 10 min at 37◦C, subjected the samples to a 10-30 % sucrose density gradient centrifugation, and analyzed the gradient fractions by radioactivity counting.

### CryoEM sample preparation and data acquisition

Aliquots of 3μl of assembled ribosome complexes at concentration range of 250-350 nM were incubated for 30 seconds on glow-discharged holey gold grids (*68*)(UltrAuFoil R1.2/1.3). Grids were blotted for 2.5s and flash cooled in liquid ethane using an FEI Vitrobot. Grids were transferred to an FEI Titan Krios microscope equipped with an energy filter (slits aperture 20eV) and a Gatan K2 detector operated at 300 kV. Data was recorded in counting mode at a magnification of 130,000 corresponding to a calibrated pixel size of 1.08 Å. Defocus values ranged from 1-3.6 μm. Images were recorded in automatic mode using the Leginon (*69*) and APPION (*70*) software and frames were aligned using the Relion3 (*47*) implementation of the Motioncor2 algorithm (*71*).

### Image processing and structure determination

Contrast transfer function parameters were estimated using GCTF (*72*) and particle picking was performed using GAUTOMACH without the use of templates and with a diameter value of 260 pixels. All 2D and 3D classifications and refinements were performed using RELION. An initial 2D classification with a 4 times binned dataset identified all ribosome particles. A consensus reconstruction with all 40S particles was computed using the AutoRefine tool of RELION. Next, 3D classification without alignment (four classes, T parameter 4) identified a class with unambiguous density for eIF3. This class was independently refined, and further masked classification allowed the identification of two subclasses distinguishable by different degree of 40S head swiveling and presence or absence of eIF3d density. Final refinements with unbinned data for the classes selected yielded high resolution maps with density features in agreement with the reported resolution. Local resolution was computed with RESMAP (*73*).

### Model building and refinement

Models for the mammalian 40S and eIF3 docked into the maps using CHIMERA (*74*) and COOT (*75*) was used to manually adjust these initial models. 5’-UTR-IRES was built manually using COOT. An initial round of refinement was performed in Phenix using real-space refinement (*76*) with secondary structure restraints and a final step of reciprocal-space refinement with REFMAC (*77*). The fit of the model to the map density was quantified using FSCaverage and Cref and model-to-maps over-fitting tests were performed following standard protocols in the field (*78, 79*).

## Supplementary figures

## Supplementary figures

**Supplementary Figure S1.**
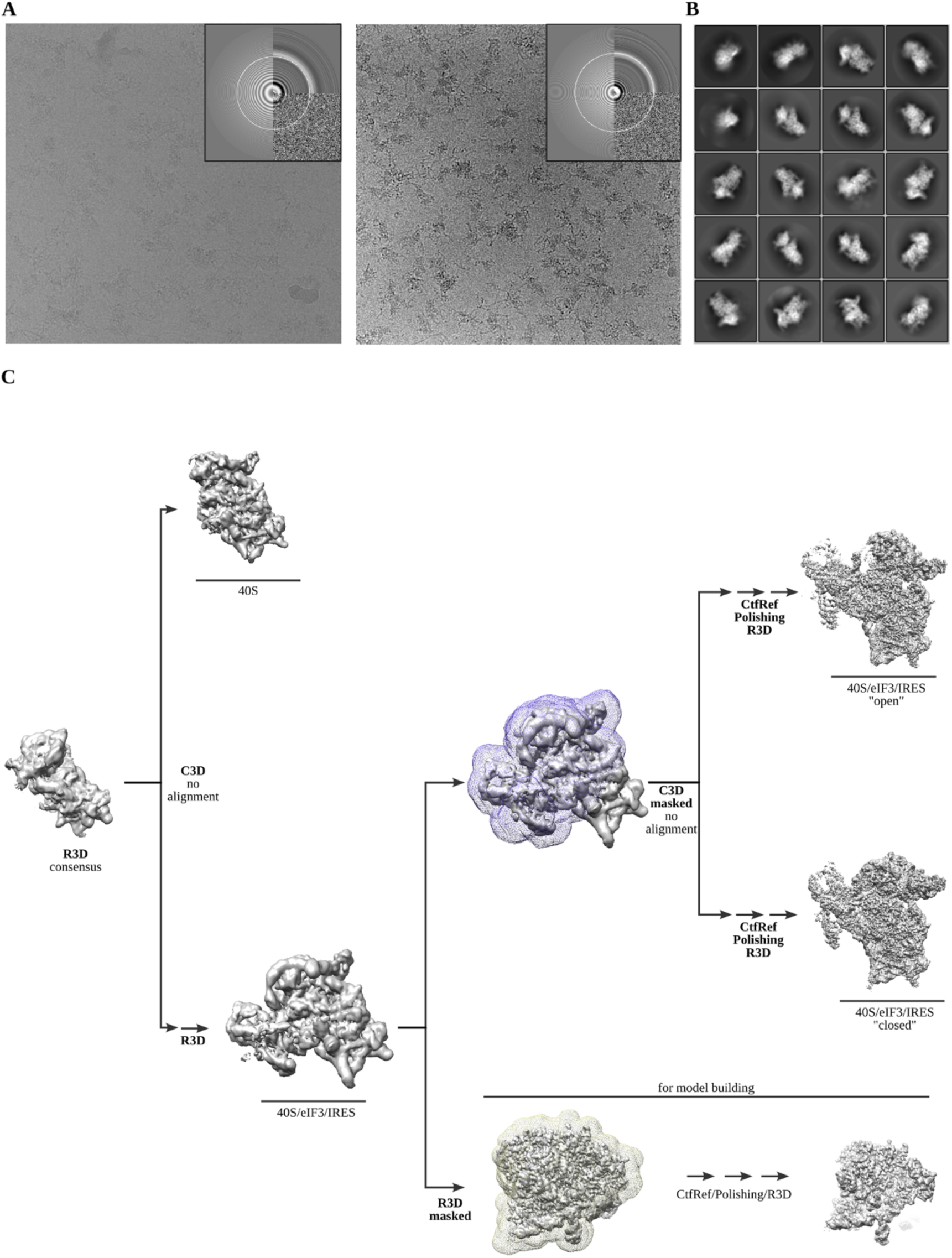
Cryo-EM representative images and classification workflow. **A,** Two examples of aligned micrographs used for image processing. Data collection in thick ice was instrumental to avoid complex disassembly and preferential orientation. **B**, Representative reference-free 2D averages. **C**, Classification scheme followed to identify the two classes described.

**Supplementary Figure S2.**
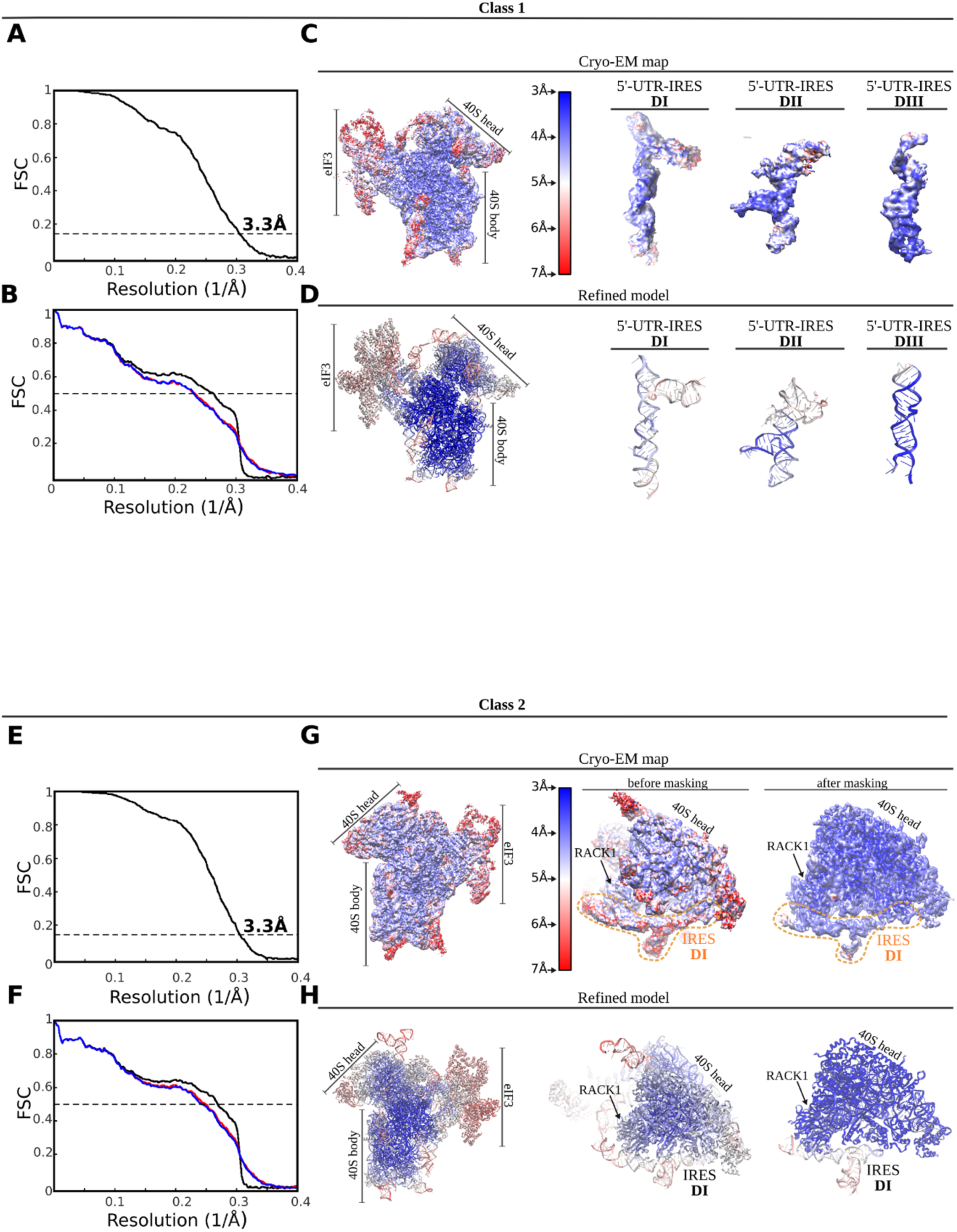
FSC correlation curves, local resolution and model validation. **Top, class-1:** **A,** Gold-Standard <u>F</u>ourier <u>S</u>hell <u>C</u>orrelation (FSC) curves between half-maps independently refined in Relion3. Global resolution by the 0.143 cutoff criterium was estimated to be 3.3Å. **B,** FSC between the final refined model and final map (black curve). Absence of model overfitting is demonstrated by the overlapping of FSC curves between half-map 1 (included in the refinement, blue) and the model and half-map 2 (not included in the refinement, red). **C**, Unsharpened map colored according to local resolution as computed by ResMap. On the right, detailed views for the three 5’-UTR-IRES domains. **D**, Refined model colored according to estimated B-factors computed by Refmac. **Bottom, class-2.** **E,** Gold-Standard <u>F</u>ourier <u>S</u>hell <u>C</u>orrelation (FSC) curves between half-maps independently refined in Relion3. Global resolution by the 0.143 cutoff criterium was estimated to be 3.3Å. **F,** FSC between the final refined model and final map (black curve). Absence of model overfitting is demonstrated by the overlapping of FSC curves between half-map 1 (included in the refinement, blue) and the model and half-map 2 (not included in the refinement, red). **G**, Unsharpened map colored according to local resolution as computed by ResMap. On the right, detailed views for the 40S head before and after masked refinements. **H**, Refined model colored according to estimated B-factors computed by Refmac.

**Supplementary Figure S3.**
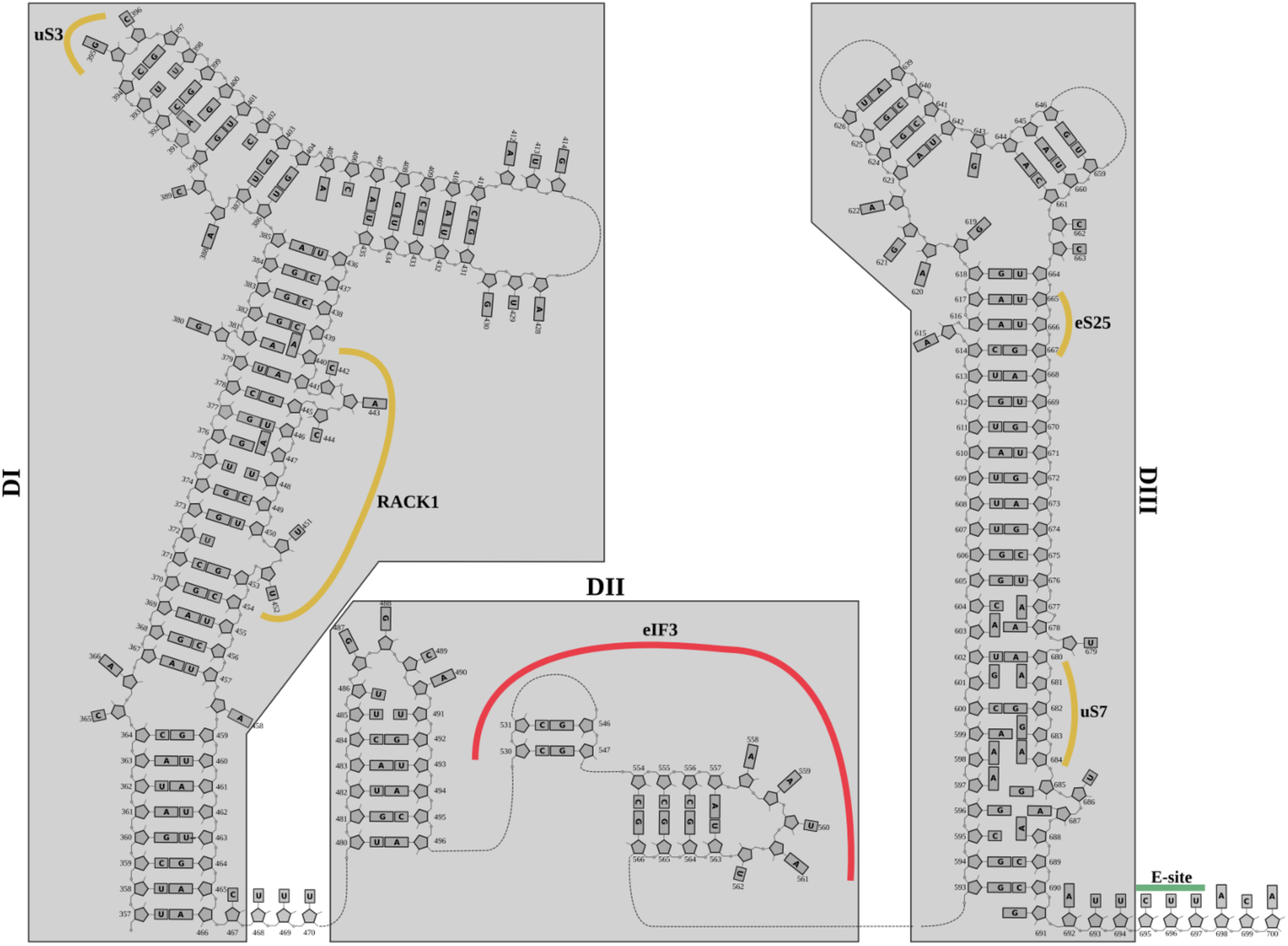
Structurally derived secondary structure diagram for 5’-UTR-IRES. Secondary structure diagram for the 5’-UTR-CrPV IRES derived from the cryo-EM structure.

Cryo-EM data collection, refinement and validation statistics.

|  | Class-1<br>(open)<br>(EMDB-XXX)<br>(PDB XXXX) | Class-2<br>(closed)<br>(EMDB-YYYY)<br>(PDB YYYY) |
| --- | --- | --- |
| **Data collection and processing** |  |  |
| Magnification |  | 1300,000 |
| Voltage (kV) |  | 300 |
| Electron exposure (e-/Å<sup>2</sup>) |  | 59.55 |
| Defocus range (μm) |  | -1/-3 |
| Pixel size (Å) |  | 1.06 |
| Symmetry imposed |  | C1 |
| Initial particle images<br>(no.) |  | 915,647 |
| Final particle images<br>(no.) | 14,257 | 23,444 |
| Map resolution (Å) | 3.3 | 3.3 |
| FSC threshold | 0.143 | 0.143 |
| Map resolution range (Å) | 3-8 | 3-8 |
| **Refinement** |  |  |
| Initial model used (PDB<br>code) | 5A2Q | 5A2Q |
| Model resolution (Å) | 3.6 | 3.6 |
| FSC threshold | 0.5 | 0.5 |
| Model resolution range<br>(Å) | 3.3-8 | 3.3-8 |
| Map sharpening *B* factor<br>(Å<sup>2</sup>) | -31.94 | -43.41 |
| Model composition |  |  |
| Non-hydrogen atoms | 28,877 | 28,877 |
| Protein residues | 28,877 | 28,877 |
| Ligands | - | - |
| *B* factors (Å<sup>2</sup>) |  |  |
| Protein | 92.47 | 96.5 |
| Ligand | - | - |
| R.m.s. deviations |  |  |
| Bond lengths (Å) | 0.014 | 0.014 |
| Bond angles (°) | 1.77 | 1.78 |
| Validation |  |  |
| MolProbity score | 2.34 | 1.81 |
| Clashscore | 6.89 | 1.82 |
| Poor rotamers (%) | 18.62 | 18.6 |
| Ramachandran plot |  |  |
| Favored (%) | 81.38 | 81.41 |
| Allowed (%) | 96.09 | 96.55 |
| Disallowed (%) | 3.91 | 3.45 |

